# Multi-protein chimeric antigens, a novel combined approach for efficiently targeting and blocking the blood stage of *Plasmodium falciparum*

**DOI:** 10.1101/2023.11.22.568251

**Authors:** Bhagyashree Deshmukh, Dhruv Khatri, Sanjay Kumar Kochar, Chaitanya Athale, Krishanpal Karmodiya

## Abstract

*Plasmodium falciparum*-induced malaria remains a fatal disease affecting millions of people worldwide. Mainly, the blood stage of malaria is highly pathogenic and symptomatic, rapidly damaging the host organs and occasionally leading to death. Currently, no vaccines are approved for use against the blood stage of malaria. Canonical vaccines in the past have selected the most immunodominant or essential protein to block the growth of the parasite. This strategy works efficiently for low-complexity organisms such as viruses and a few bacteria but has not shown promising results for a malaria vaccine. *Plasmodium* has a complex life cycle and vaccine candidates especially during blood stage are ineffective due to multiple gene families showing redundancy, immune evasion, and insufficient antibody titer. Herein, we demonstrate a novel strategy of combining multiple antigens from the blood stage of *Plasmodium falciparum* using only the most immunodominant peptide sequences as a way of tackling polymorphism and redundancy. We created three chimeric antigens targeting eight PfEMP1 proteins (chimeric varB) and eight merozoite surface proteins (chimeric MSP and InvP) by selecting and stitching B-cell epitopes. Our chimeric constructs show naturally circulating antibodies against individual peptides using epitope-mapping microarray as well as entire proteins in malaria-infected patients. We demonstrate that anti-varB antibodies are neutralizing in nature and significantly reduce the cytoadhesion on an organ-on-chip system with a microfluidic device mimicking physiological conditions. We have applied a Deep Learning based method to quantify the number of adhered RBCs under fluidic conditions that is used to study cytoadhesion. Furthermore, the anti-MSP and InvP antibodies show complete growth inhibition in a single cycle at a combined concentration of 0.13 mg/ml. Overall, our results show that a combination of antigenic peptides from multiple antigens can function as a next-generation vaccine and effectively block the blood stage by reducing cytoadhesion and inhibiting the parasite growth.

## Introduction

Malaria remains a public health threat in developing countries despite decades of efforts in control and eradication. The burden of malaria in 2021 included 247 million cases and 625,000 deaths worldwide (*1*). The WHO has approved the RTS,S/AS01 vaccine for malaria, targeting the pre-erythrocytic stage. However, its efficacy has been sub-optimal, with the effect waning over time (*2*). Another vaccine recently approved, R21/Matrix-M, has shown slightly better efficacy but has limitation in targeting only the pre-erythrocytic stage (*3*). The clinical manifestation of malaria is observed during its blood stage, wherein multiple parasite-derived proteins lead to the severity of symptoms, complications, such as cytoadhesion and rosetting, and increased parasite density by repeated multiplication cycles (*4*). Despite this highly damaging stage, our only intervention against the blood stage is using antimalarial, such as artemisinin combination therapy (ACT), to reduce the parasitic load (*5, 6*). However, the recent emergence of drug resistance and other complications during the blood stage calls for effective interventions alongside antimalarial drugs (*7–10*).

Generating vaccine against the blood stage has always been challenging due to the complexity of the stage including high virulence, faster multiplication, lower antibody yield and many target candidates failing during phase trails (*11, 12*). The targeting of blood stage till now has primarily focused on finding the best target protein to clear disease progression and alleviate severity efficiently. However, redundancy in the pathways and the ability of the parasite to evade the host immune system makes it nearly impossible to rely on just one protein or one stage as an effective vaccine target (*11*). It has been well established that with multiple exposures, the breadth of immune response and circulating antibodies against various antigens can reach a threshold where infection remains asymptomatic or clears over time. This phase is achieved when multiple immunodominant proteins are exposed to the immune system repeatedly and substantial amount of neutralizing antibodies are generated in the host body (*13*). These antibodies if passively transferred to a naive infection patient is known to reduce the disease severity significantly (*14–16*). Collectively, these observations strongly suggest that vaccination against the blood stage is possible and the induced antibodies can be protective in nature.

The blood stage of infection starts with a merozoite entering into an erythrocyte via a multi-step mechanism. The merozoite initially attaches with the RBC surface through its vast MSP family proteins. This initial attachment keeps the merozoite latched onto the surface. It is followed by reorientation process wherein the apical end of the merozoite faces the RBC membrane. A parasite-protein driven tight junction is made that serves as a bridge for merozoite entry and using actin-myosin movement, the merozoite enters into the RBC leaving a protein coat behind. Once inside, the *Plasmodium falciparum* takes 48 hours to divide and mature into daughter merozoites that further keep the cycle perpetuating (*17–19*). During this stage, the parasite remains hidden in the RBC but also expresses certain virulence proteins that are exported to the surface of the RBC and help them adhere to the endothelial lining of the blood vessel. This allows the parasite invaded RBC to remain elusive to the immune system and avoid the splenic clearance. This cytoadhesion and sequestration is majorly caused by the parasite protein family *Plasmodium falciparum* erythrocyte membrane protein 1 (PfEMP1) that has ∼60 genes encoding them, has multi-domain structure with variations, shows mutually exclusive expression and contributes to the severity of the disease (*20, 21*). This sequestration of infected RBCs (iRBCs) in vital organs such as brain, kidney, lungs etc can make the infection fatal (*22–25*). Malaria-exposed individuals do show presence of anti-PfEMP1 antibodies which have shown to be neutralizing and protective in nature (*26–30*). Overall, the PfEMP1 proteins and the merozoite surface proteins together make ideal candidates that have immunodominancy and can be targeted to reduce the disease burden and severity.

Seroepidemiological studies have shown that the presence of IgG-specific functional antibodies against merozoite antigens, specifically AMA-1, MSP-2, and MSP-3 in combination, correlated with greater protection against clinical malaria (*31*). The presence of functional antibodies against virulent family proteins such as PfEMP1s protects against severity and complications (*26, 28–30*). Protein candidates, such as RH5, EBA-175, and GLURP, are well-considered vaccine targets, with RH5 currently being in phase trials (*32–35*). However, a recombinant single protein target has limitations in inducing a sufficient antibody titer and level of security. Compared with the canonical single-protein approach, multiple studies have focused on generating next-generation targets, including highly immunogenic combinations of multiple proteins that can deliver the desired effects. Research in generating chimeric epitope-based vaccines against ascariasis, hepatitis C, Leishmaniasis and Toxoplasmosis have shown promising result (*36–39*). However, this approach has not been systematically explored in case of malaria vaccine design.

In this study, we have used a novel strategy of combining the antigenic B-cell recognizable peptide stretches from multiple proteins as a way of increasing the target breadth. We selected antigens based on their surface-localization, expression window during blood stage, known immunodominancy, polymorphic nature and evidence of protective antibodies in natural infections. We constructed, expressed, and validated three chimeric antigens targeting two different blood stages of *Plasmodium falciparum*. For construction, we used the highest-scoring B-cell epitopes of a total of eight PfEMP1 subgroup B (chimeric varB), four merozoite surface proteins (chimeric MSP), and four merozoite invasion-specific proteins (chimeric InvP). We observed that naturally circulating antibodies recognize the epitopes used in construction, with significantly higher titers in the severe form of malaria. We show that antibodies specific to chimeric varB reduce cytoadhesion under static and fluidic conditions. Antibodies specific to chimeric MSP and InvP constructs showed complete growth inhibition with additive effects. Thus, this study demonstrates that the multi-protein chimeric antigens efficiently target and block the blood Stage of *Plasmodium falciparum*.

## Results

### *In-silico* epitope prediction and generation of chimeric antigens

The severity of malaria is usually caused in the blood stage by parasite-derived proteins exported to the RBC surface exhibiting the ability to evade immune recognition and avoid splenic clearance. One such family of proteins is PfEMP1 which interact with endothelial receptors of the blood lining and cause sequestration of the iRBC leading to high inflammation and microcapillary obstruction (*21*). Interestingly, despite being surface-exposed, immuno-dominant and show presence of neutralizing antibody repertoire upon multiple exposure, this family of proteins are not considered as ideal vaccine candidates due to a large number of genes per family, mutually exclusive expression, and their ability to switch from one gene to another within a single host (*40, 41*). To test our hypothesis, if multi-protein chimeric antigens are better target than a single protein, we selected clinically relevant PfEMP1 family of proteins involved in cytoadhesion and sequestration (Figure 1A) and merozoite surface proteins that aid in the invasion process (Figure 1B). We re-analyzed the RNA-sequencing data of 1027 patient samples (published by Mok et al., Science, 2015) to identify the highly expressed PfEMP1 candidate genes (*42*). In the uncomplicated form of *falciparum*-malaria infection, PfEMP1 subgroup B was dominantly expressed compared to the other subgroups (Figure 1A). As proof of principle, we picked eight proteins (PF3D7_1200100, PF3D7_0500100, PF3D7_1300100, PF3D7_0324900, PF3D7_1100100, PF3D7_0937800, PF3D7_0300100, and PF3D7_0400100) belonging to group B, which also had dominant expression in the lab strain 3D7 (Supplementary Fig 1A). This overlap of gene expression made it feasible to use the lab strain for further downstream experiments. The protein sequences of selected candidates were taken from PlasmoDB and subjected to the Immune Epitope Database-Analysis Resource (IEDB-AR) for predicting the antigenicity of the peptides, and a total of six parameters were considered for scoring the antigenicity (details provided in method section) (*43*). A total of 29 peptides originating from the functional domains of a length of 25 amino acids were stitched together to create the chimera varB (Supplementary Table 1; Sheet 1). The protein sequence was back converted into a codon-optimized DNA sequence, cloned into pET28a+ plasmid with N-terminal 6X histidine tag, and expressed in the *E. coli* system under the control of the T7 promoter and used for further assays (Figure 1C).

**Figure 1.**
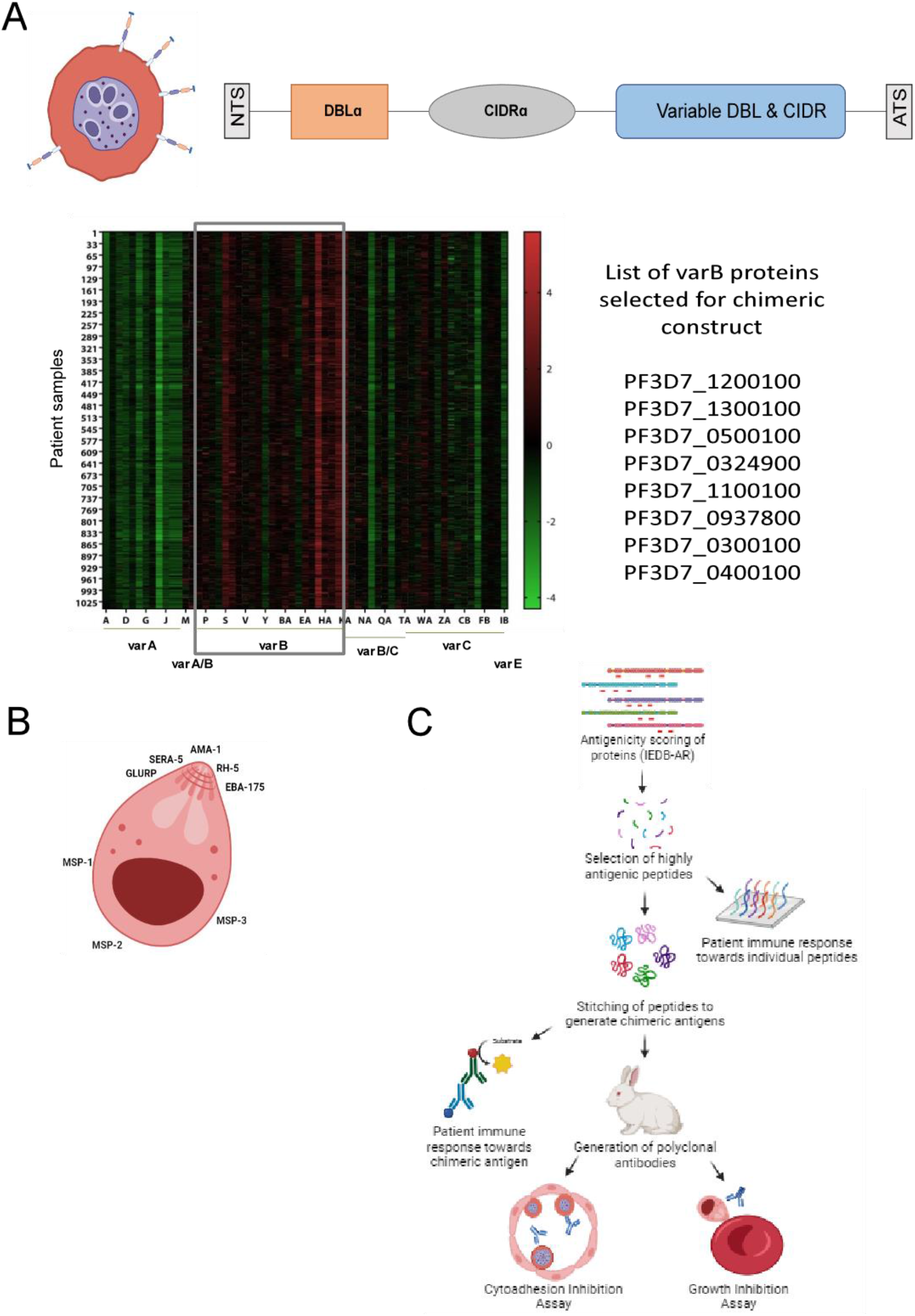
Successful construction, expression and validation of chimeric antigens. **(A)** Structural domains present in PfEMP1 proteins and a heatmap demonstrating higher expression of varB subtype in 1027 patient samples (Re-analyzed from Mok et al., 2015) along with the list of eight varB proteins selected for generation of chimeric varB antigen. **(B)** Structure of merozoite and the location of proteins selected for the generation of chimeric MSP and InvP construct. **(C)** The workflow depicting the construction of chimeric varB, MSP, and InvP antigen.

Similarly, we selected the invasion and merozoite-specific proteins (MSPs) which are known to be immunodominant and already validated as potential vaccine candidates individually (*33–35, 44–46*). For constructing chimeric MSP, we predicted highly antigenic peptide stretches of 25 amino acids from AMA-1 (PF3D7_1133400), MSP-1 (PF3D7_0930300), MSP-2 (PF3D7_0206800), and MSP-3 (PF3D7_1035400), and stitched them together. A chimeric InvP (invasion proteins) construct was generated by stitching the highly antigenic peptides from EBA-175 (PF3D7_0731500), SERA-5 (PF3D7_0207600), RH-5 (PF3D7_0424100), and GLURP (PF3D7_1035300) (Figure 1B). The three chimeric constructs, i.e., chimeric varB, MSP, and InvP, with a final average antigenicity score above 0.5 (Supplementary Table 1; Sheet 2 and 3 respectively) were expressed, purified individually, and validated using mass-spectrometry (Supplementary Fig 1B and 1C). Thus, we successful constructed chimeric antigens by stitching highly immunodominant peptides, cloned and expressed them in the bacterial expression system. These three constructs targeting eight PfEMP1 proteins expressed on surface of iRBCs (Chimeric varB) and eight merozoite proteins present on merozoite surface (chimeric MSP and InvP) cover the entire blood stage of *Plasmodium falciparum* with parasite targeted while being inside as well as outside of the host RBCs.

### Naturally circulating antibodies show reactivity against chimeric antigens

In order to know if the in-silico identified epitopes are exposed to human immune system and if there are circulating antibodies against them, we performed an ELISA assay using serum samples from malaria-infected individuals. The malaria-infected individuals show significant reactivity in comparison with healthy controls against chimeric varB, MSP, and InvP (Figure 2A). Moreover, the severe malaria samples react significantly more toward the chimeric MSP and InvP construct (Figure 2B). One of the 30 severe malaria patient samples had a significantly higher response toward the chimeric varB construct. This could be owing to the diversity of PfEMP1 protein expression during infection. Through this assay, we observed that the antigen predicted using in-silico analysis are exposed to immune system in patient samples during natural infection and are being recognized by the naturally circulating IgG antibodies. We also performed Immunoprecipitation followed by mass-spectrometry of parasite lysate using serum samples of malaria-infected individuals of Indian cohort. The selected eight proteins were also exposed in the patients during natural infection further confirming our selected chimeric antigens. The heterogeneity of expression of var proteins and antibody titer was clearly seen within these candidates (Supplementary Figure 1D). It is well known that presence of cytophilic IgG antibodies against the proteins considered in the generation of chimeric antigens are known to be protective in nature and correlate positively with the disease survival (*47, 48*). Hence the IgG response against the designed antigens is also very likely to be highly protective against the infection.

**Figure 2.**
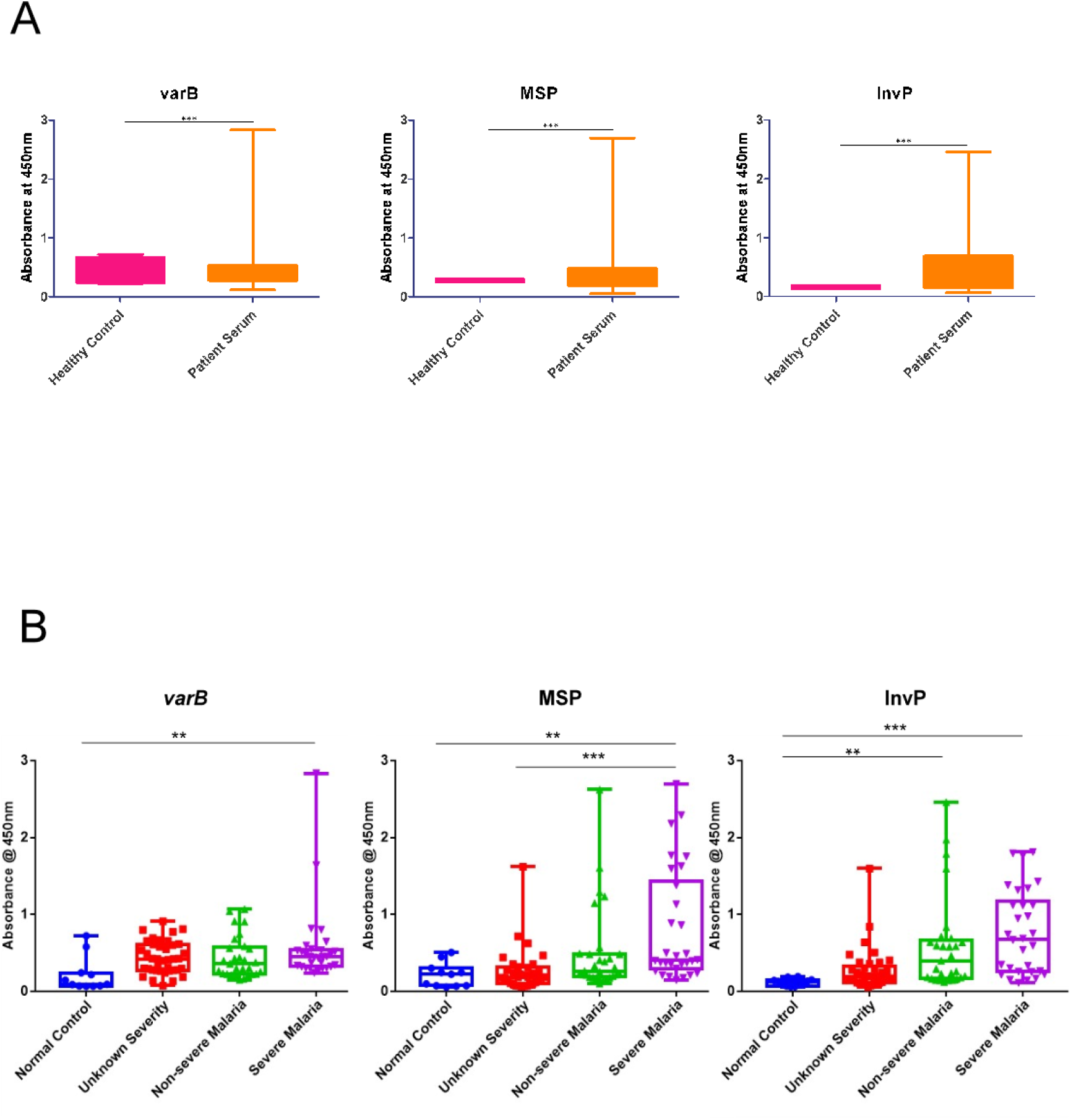
Naturally circulating antibodies recognise the chimeric antigens. **(A)** Plots depicting mean OD for IgG reactivity of malaria-infected individuals in comparison with healthy controls against chimeric varB (left), MSP (center), and InvP (right). (**B)** The IgG reactivity of malaria-infected individuals based on the severity of malaria. One-way ANOVA, multiple comparison test, P<0.05.

### Individual peptides used in chimeric antigens show heterogeneity in immune response indicating polymorphism in antigen exposure

Further to know if the naturally circulating antibodies against chimeric antigens are a result of a combination of peptides used and to observe the polymorphism in antigen expression, we performed peptide-mapping microarray, where we looked at the human immune response to individual peptides used in the chimeric construct. This allows an epitope-level resolution to the antibody responses against the peptides and helps in better understanding of heterogeneity and unique antibody responses against each peptide used. The microarray design included peptides in duplicate tethered to a slide (Epitope mapping by PEPperPRINT, GmbH, Germany) that can be readily probed using antibodies. The peptides selected in constructing chimeric varB, MSP, and InvP, along with HA and polio peptides as control, were synthesized and imprinted. We incubated the imprinted arrays with patient serum samples to observe the peptide-specific IgG and IgM responses. We observed a wide range of patient immune responses against peptides selected in all three chimeric antigens correlated with their respective ELISA values (Supplementary Figure 2). Every peptide used in constructing chimeric varB had IgG reactivity, and this response was maximum towards peptides belonging to varB proteins PF3D7_1300100 and PF3D7_1200100 (Figure 3A). The average IgG response of the peptides used per protein shows a clear heterogeneity in the expression of varB genes. Across 19 patient samples the level of immune response to a particular varB subtype protein varied greatly indicating that not all the PfEMP1 are expressed in every individual and hence a combined approach is more relevant in case of targeting virulence multi family proteins (Figure 3A). The chimeric MSP construct showed uniform reactivity across the peptides selected (Figure 3B). Four patient samples showed reactivity against 75% of the peptides used, while a broad range of reactivity was seen against the peptides belonging to AMA-1 and MSP-3. The average IgG response per protein showed a trend where AMA-1, MSP-1 and MSP-2 showed highest IgG peaks in different patient samples with no overlap. While MSP-3 antibodies were present in lower titres but were present across all 19 patient samples (Figure 3B). The peptides of the chimeric InvP construct had a steady IgG reactivity across. The peptides belonging to RH-5 and SERA-5 proteins showed maximum reactivity in 8 patients out of 19 (Figure 3C). We clearly see one dominating protein being recognized in the patient samples after analysis of average IgG per protein approach. The patient samples that show highest SERA-5 antibodies show a lower recognition by other proteins. Similarly, where the expression of EBA-175 or RH-5 was maximum the IgG response to other proteins was lower (Figure 3C). The peptide-mapping for all chimeric antigens shows that every host produces a varied levels of IgG responses against different proteins. This result clearly indicate why single protein vaccine works only in a subset of population and fails to elicit protective antibody response in all individuals. To achieve a threshold of protective immune response considering multiple antigens is a necessity.

**Figure 3.**
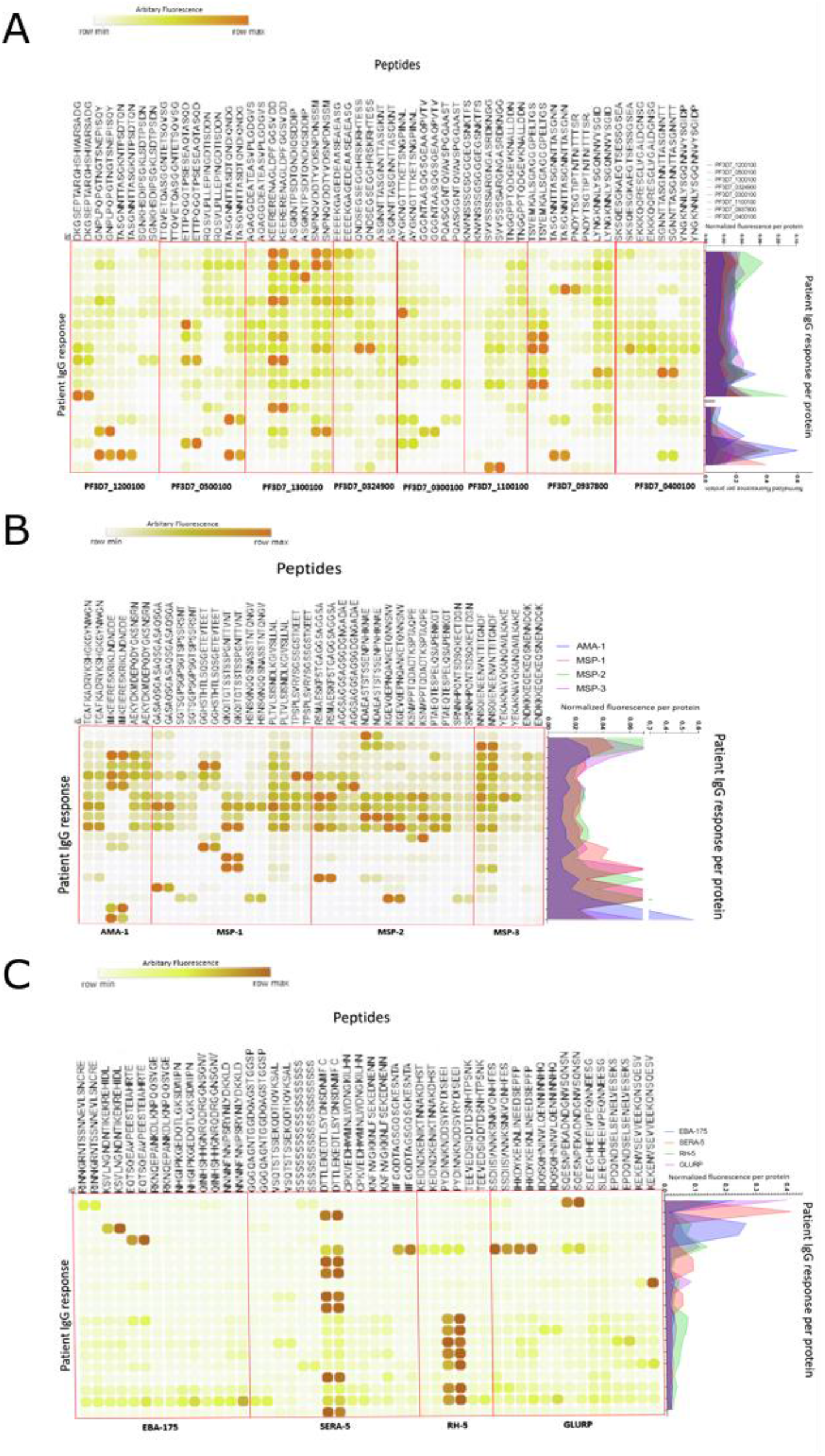
Epitope-mapping microarray shows unique IgG response towards each peptide. **(A)** Represents the IgG response of 19 malaria-infected individuals against each peptide used in chimeric varB antigen. A bar graph to the right depicts the ELISA response of the respective serum sample to the entire chimeric varB antigen. **(B)** IgG response of 19 malaria-infected patient samples towards each peptide used in chimeric MSP antigen and ELISA reactivity towards complete construct. **(C)** IgG reactivity of malaria-infected patient samples towards peptides of chimeric InvP with correlating ELISA value for the entire construct.

We also looked for IgM responses against the antigens and observed a trend similar to that of IgG (Supplementary Figure 2). The IgM response in chimeric varB was higher in peptides belonging to PF3D7_1300100, PF3D7_1200100, and PF3D7_0937800 (Supplementary Figure 2A). The heterogeneity expected in the exposure to the PfEMP1 proteins was seen across IgG and IgM response to chimeric varB construct. Certain patient samples had antibodies present against all of the peptides used in the construct while others show very protein specific response. IgM response against chimeric MSP showed higher breadth towards AMA-1 peptide followed by MSP-1. Around 10 samples out of 19 had strong IgM response against all peptides used (Supplementary Figure 2B). For chimeric InvP the IgM response was higher in peptides of RH-5, one peptide in SERA-5 and GLURP. All the proteins selected for the chimeric construction are known for their polymorphic nature and hence the variation in immune response was as expected. Thus, a comprehensive analysis revealed extensive IgG and IgM responses in patient samples targeting the full spectrum of peptides incorporated into chimeric antigens. This underscores the imperative to shift from singular protein candidates to multi-protein chimeric antigens when developing vaccines. This necessity arises from the intrinsic polymorphic diversity inherent in antigens, showcasing the essentiality of a broader immune response for effective vaccination strategies.

### Chimeric varB-specific antibodies reduce the rosette formations and cytoadherence

All three chimeric antigens show presence of immune response across patient samples in the form of whole antigen as well as on individual peptide level. The presence of IgG response would intuitively mean a possible neutralizing effect. To test whether the chimeric antigen specific IgG antibodies are functional in nature, we generated a polyclonal antibody response against them in rabbits. We tested the anti-chimeric varB antibodies for their ability to inhibit the rosetting and cytoadhesion in culture conditions. While the anti-chimeric MSP and InvP specific antibodies were used to determine the invasion inhibition potential.

The bleed collected for anti-chimeric varB antibody was subjected to IgG enrichment using affinity protein G beads. To confirm the specificity of the generated antibodies, we tested them on peptide-microarray containing all the peptides used in construction of chimeric varB. The rabbit antibodies had a broad range of reactivity against the peptides, with peptide belonging to protein PF3D7_1200100 having the highest titer of rabbit IgG followed by PF3D7_1300100, PF3D7_0324900, and PF3D7_0400100 (Supplementary Figure 3A). Once the breadth and specificity of the immune response was confirmed, we performed rosetting and cytoadherence-inhibition assays to test the neutralizing potential of the chimeric varB-specific antibodies. The rosetting assay was done by wet mounting followed by manual counting as well as flow cytometry-based methods as described earlier (*49*). In 3D7 culture, we observed the formation of rosettes defined by two or more uninfected RBCs binding to a common iRBC (Figure 4A). Adding chimeric varB-specific antibodies reduces the number of rosettes formed by 30% at the 0.01 mg/ml concentration as seen by FACS sorting of multiplet population (Figure 4B). We did not observe complete inhibition of rosette formation, which can be due to the expression of PfEMP1 genes other than the eight selected in making the chimeric construct.

**Figure 4.**
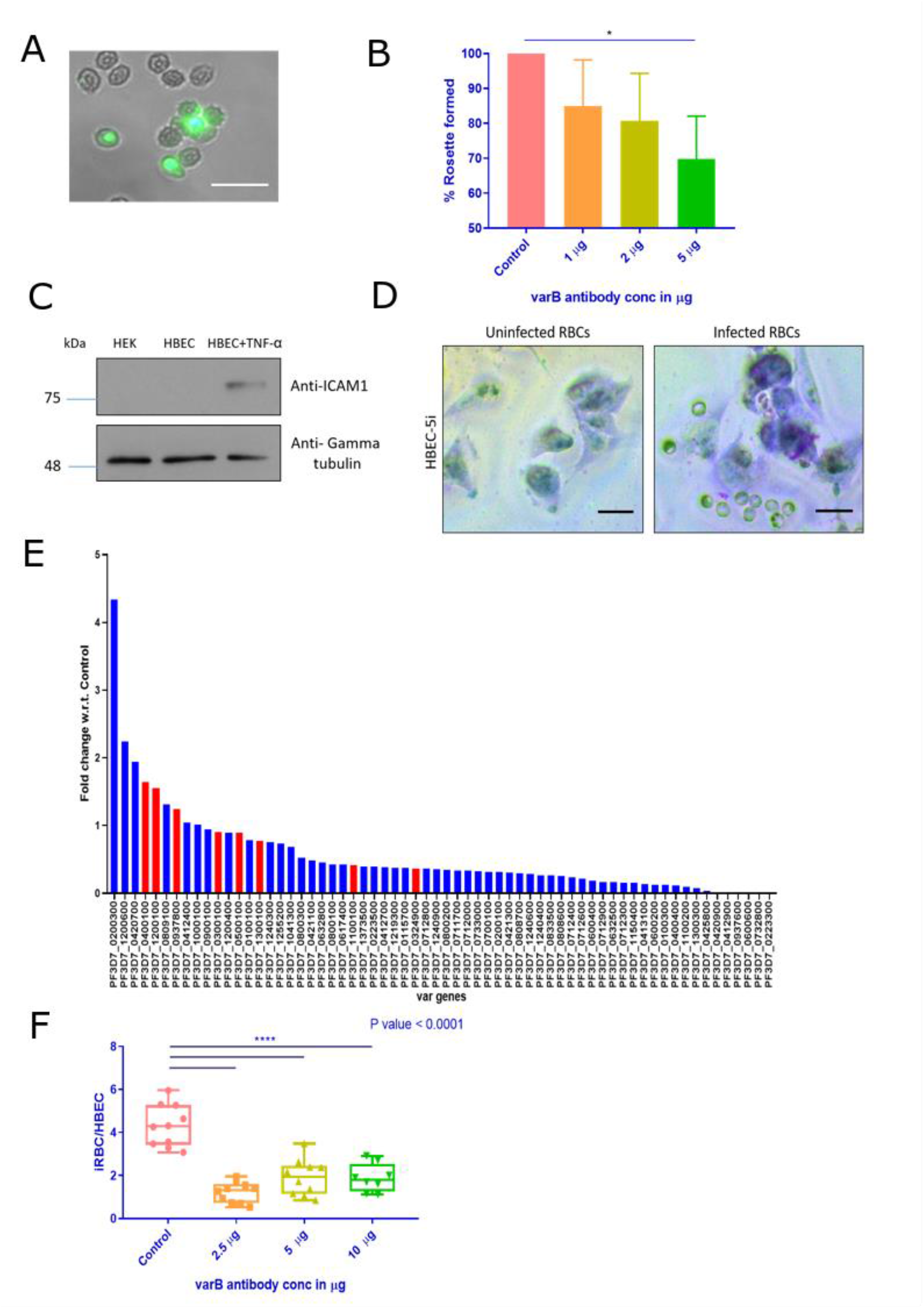
Chimeric varB specific antibodies show neutralizing effect. **(A)** A rosette structure of central iRBC stained with acridine orange surrounded by uninfected RBCs. **(B)** A plot showing a reduction in the % of rosette formed upon the addition of chimeric varB specific antibodies. **(C)** A western blot image for ICAM-1 levels in HBEC-5i cells with and without TNF-ɑ treatment. Actin is shown for loading control. **(D)** A microscopic image of Giemsa stained iRBCs showing cytoadherence to primed HBEC-5i cells. **(E)** A plot of the PfEMP1 gene-specific Transcript per million (TPM) values of HBEC-5i enriched iRBCs. The red mark shows the levels of varB-specific genes used to generate chimeric varB antigen. **(F)** A plot showing iRBCs to HBEC-5i cell ratio of static cytoadherence assay under control and varB-specific antibody treatment condition.

Apart from rosetting, the PfEMP1 proteins are well known for their binding affinity with human endothelial receptors and causing sequestration of iRBCs in deep capillaries leading to severe malaria (*20, 24, 50, 51*). Population living in endemic regions having multiple exposure to the infection are known to have functional circulating antibodies against PfEMP1 that corelated with reduced severity associated with cytoadhesion and better survival (*26, 28–30, 50, 52*). Since we are expecting the anti-chimeric varB antibodies to function on similar basis, we tested their neutralizing efficiency using a human endothelial cell line. We have used human brain endothelial cells (HBEC-5i) primed with TNF-ɑ as our model for testing the cytoadhesion (*53*). These primed endothelial cells show expression of ICAM-1 receptor which is well known for binding to PfEMP1 especially in cerebral malaria cases (Figure 4C). We observed a strong affinity of iRBCs to the primed HBEC-5i cells whereas the healthy RBCs served as negative control for cytoadhesion (Figure 4D). Since a parasite population in a cell line is known to have a diverse expression of PfEMP1 genes, it is vital to first confirm the role of selected PfEMP1 genes in the cytoadhesion process. To find the PfEMP1 expression in the HBEC-5i binding population, we performed RNA-sequencing of enriched cytoadhered population. As expected, the population had multiple PfEMP1 genes expressed, and the candidates selected in constructing the chimeric varB were well represented in the population (Figure 4E). We then performed the static cytoadherence inhibition assay during the mature stages of the parasite. We observed that adding chimeric varB-specific antibodies reduces the ratio of iRBCs bound per HBEC-5i cell while only IgG control had no effect (Figure 4F). We did not observe a complete inhibition of iRBCs probably due to the expression of PfEMP1 other than eight selected. However, a significant reduction in the cytoadhered RBCs was observed with the range of 2.5 μg-5 μg (0.005-0.01 mg/ml). These cytoadherence assays confirmed that the anti-chimeric varB antibodies are indeed neutralizing in nature and are capable of functionally inhibiting the cytoadhesion.

### Chimeric anti-varB antibodies significantly reduce the cytoadhesion under physiological conditions

Static assays have long been used to assess the ability of drugs, antibodies, or pharmacological agents to inhibit cytoadhesion (*54*). However, these assays fail to recapitulate the conditions in an *in-vivo* environment wherein the structure of blood capillaries, pressure, flow rate, etc., play an essential role in determining the cell-to-cell interaction (*55–57*). It is well known that depending on the receptor interaction, the iRBCs show rolling motion, tumbling, or firm adhesion phenotype under dynamic flow conditions (*56*). We combined an organ-on-chip system with a microfluidic device to mimic these physiological conditions (Figure 5A). The primed HBEC-5i cells were seeded onto the capillary and allowed to adhere. The parasite culture of 5% parasitemia was diluted to 0.15% hematocrit (lower hematocrit was standardized to track RBCs in real-time) and passed through the capillary at the pressure of 25 mbar. These parameters are often found in the deep capillary beds of the organ, where a majority of iRBC adherence is found. Under these conditions, healthy RBCs do not interact with the primed endothelial cells, while iRBCs show a slow-rolling phenotype and firm cytoadhesion (Supplementary Video 1). We build an analysis pipeline for detecting and counting iRBCs (Materials and Methods section). Each 1-minute video captured was divided into frames and subjected to analysis for counting the bound iRBCs per frame with a confidence of 95.75% (Figure 5B). For the quantification of RBCs, initially intensity peaks were detected from images (Supplementary Figure 3B) using a previously developed image detection pipeline for label-free detection (*58*). This was followed by the criterion of stationary vs. moving RBC and convolution network for selecting RBCs that are in-focus (Supplementary Figure 3C). We pre-incubated the enriched chimeric varB antibodies with the culture for 10 minutes at the concentration of 0.02 mg/ml and 0.04 mg/ml before passing it through the capillary. We observed a significant decrease (about 90%) in the number of bound iRBCs (Figure 5C). The decrease was consistent across the time window, and most iRBCs moved faster without rolling on the cell surface (Supplementary Video 2). With our standardized microfluidic assay combined with the real-time analysis of bound RBCs, we further affirmed the neutralizing capability of anti-chimeric varB antibodies and show that their potency is significantly higher when assessed at the physiological conditions. The real-time analysis of bound RBCs has not been shown previously and this method can prove valuable for quantifying phenomena such as cytoadhesion rate and binding/ stationary time spent by the iRBC.

**Figure 5.**
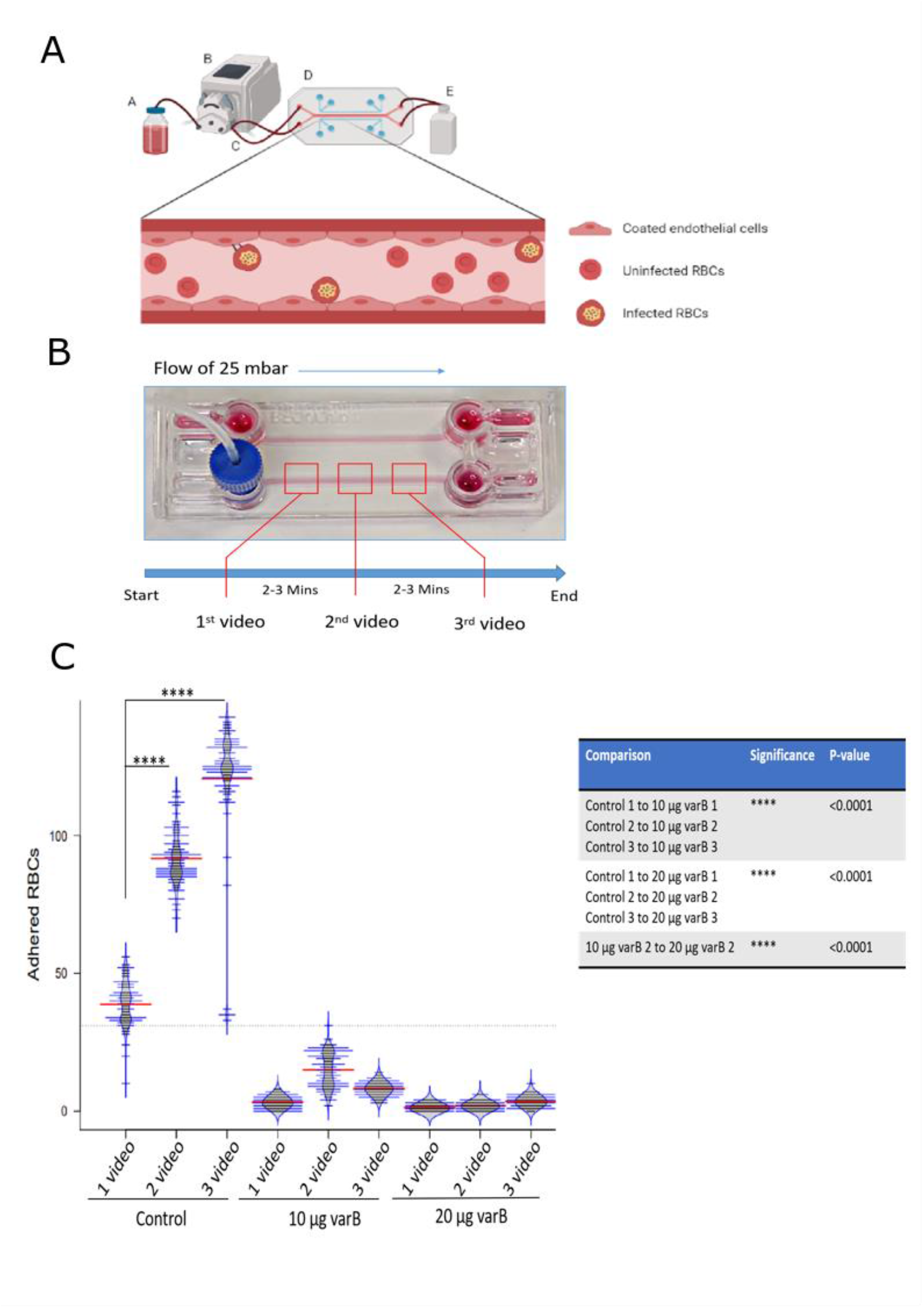
Chimeric varB antibodies show inhibition of cytoadhesion under fluidic pressure. **(A)** The design of microfluidic set up with components, A is media reservoir, B is pressure generator, C is fluidic tubings, D is flow chip and E is discard container. **(B)** Image of BeOnChip chamber (Fluigent) with the parameters used for video acquisition of in-flow adhered iRBCs. **(C)** A plot showing number of stationary RBCs per present per frame across the control and chimeric varB-specific antibody treatment condition. A table at top right corner show results of statistical analysis across all conditions. One-way ANOVA, Multiple comparison test.

### Chimeric MSP and InvP-specific antibodies inhibit parasite growth over cycles

Similar to chimeric varB, we wanted to assess the neutralizing abilities of anti-chimeric MSP and InvP antigens. These two constructs contain the peptides of eight proteins that have crucial role in merozoite’s initial attachment to the new RBC, its reorientation to align the apical side and finally form a bridge for invasion process (*17, 19*). We generated polyclonal antibodies against these constructs in rabbits and further used them for assessing their ability in blocking merozoites. The bleeds were purified using immobilized antigens as affinity columns and further validated using peptide blocking assay and immunofluorescence. Using purified antibodies, we observed multiple bands for chimeric MSP-specific antibodies precisely, three bands between size 135 to 180 (corresponding to MSP1), a band near 68 kDa (AMA1), and near 48 kDa. For chimeric InvP-specific antibodies, a strong band was observed near 135 kDa (GLURP), 100 kDa (SERA5), and multiple bands between 63 and 75 kDa. However, all the bands observed were specific to antigens as they were efficiently blocked by adding native antigens during probing (Figure 6 A and C). Since both the constructs target the merozoite surface, we performed an immunofluorescence assay at the mature schizont stage. We observed the signal on the surface of developing merozoites for both antibodies (Figure 6 B and D). We used these antibodies to assess their growth inhibitory potential alone and in combination. For this, we used growth inhibition assay (GIA), routinely used to quantify potential antibodies’ efficiency in blocking the progression to the next cycle (*59*).

**Figure 6.**
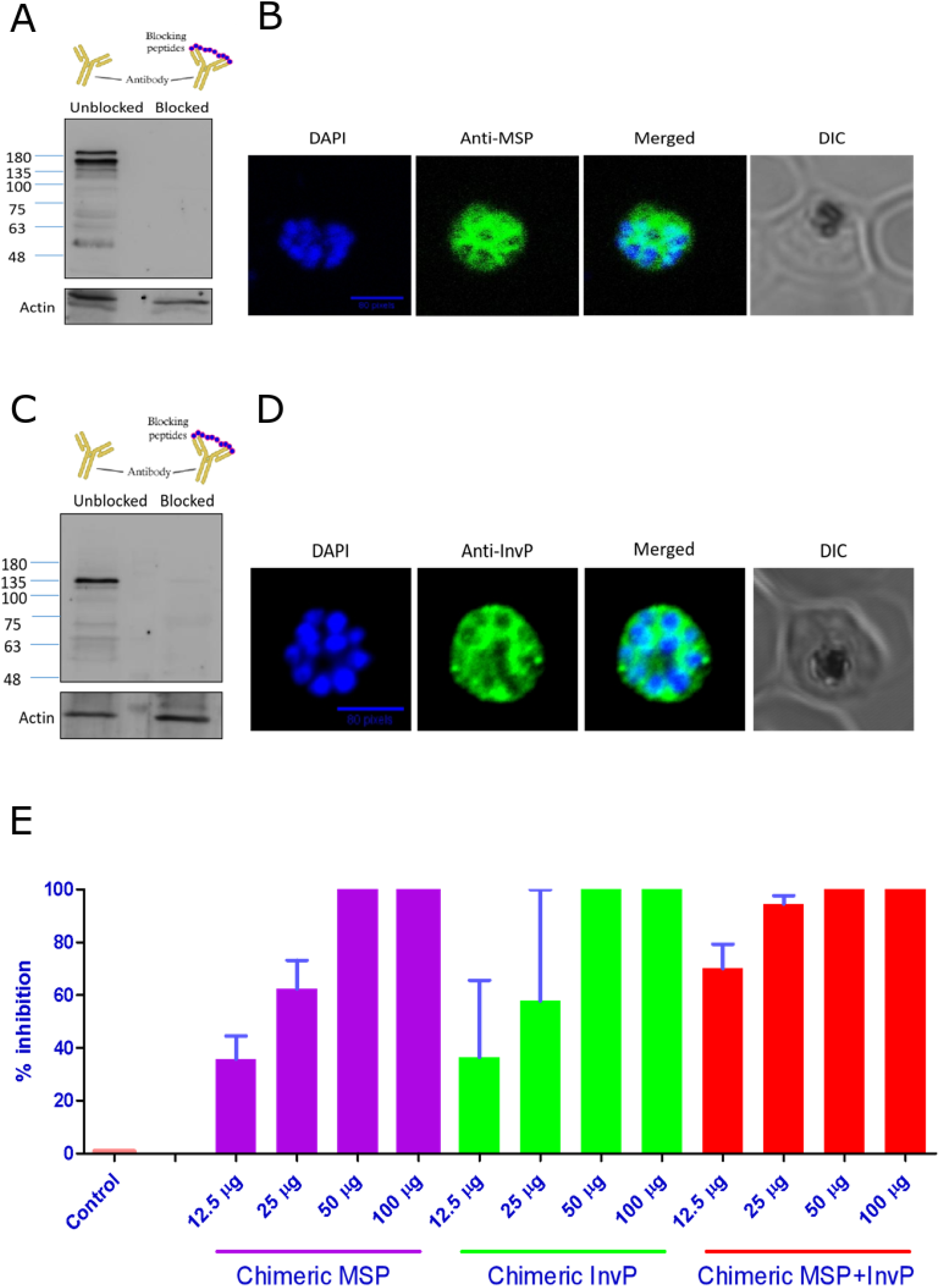
Chimeric MSP and InvP-specific antibodies show inhibition of parasite growth. **(A)** A peptide-blocking assay showing bands specific to the chimeric MSP-specific antibodies probed on 3D7 lysate**. (B)** Immunofluorescence image showing the specificity of anti-chimeric MSPantibodies recognizing the surface of matured schizont. **(C)** Peptide blocking assay showing specificity of anti-chimeric InvP antibodies probed on 3D7 lysate. **(D)** Immunofluoresence image showing binding of anti-chimeric InvP antibodies to matured schizont surface. **(E)** Single cycle growth inhibition assay showing reduction in parasitemia upon addition of chimeric MSP, InvP antibodies separate and in equivalent combination.

In single cycle GIA on the 3D7 strain, we used the antibodies in the final concentration range of 0.06 mg/ml (12.5 μg) to 0.5 mg/ml (100 μg) for both the chimeric antibodies. For chimeric MSP antibodies, 0.06 mg/ml concentration showed inhibition of around 35.7%, 0.125 mg/ml showed 62.41% inhibition, and 0.25 mg/ml concentration inhibited the parasite growth completely. For chimeric InvP antibodies, 0.06 mg/ml concentration showed inhibition of 36.45%, 0.125 mg/ml showed 57.82% inhibition, and complete inhibition at 0.25 mg/ml. When we added both the chimeric MSP and InvP antibodies in equivalent concentrations, the growth inhibition potential was amplified—the 0.06 mg/ml mix concentration yielded the inhibition of 70.1%, and 0.125 mg/ml yielded 94.55% inhibition (Figure 6E). The chimeric MSP and InvP constructs show complete inhibition with their additive effect. This shows a potent neutralizing effect of the generated antibodies in a combination.

## Discussion

Effectively targeting malaria infections remains a high-priority problem despite decade-long efforts. Research on other infectious diseases has attempted to construct a candidate with a broad antigen-targeting range. With a complex stage-specific disease, such as malaria, there is an urgent need for multi-target antigens. Here, we present a combined approach of multi-peptide, multiprotein candidate for effectively targeting the blood stage of malaria. The selected proteins are well-known immunodominant targets verified in several seroepidemiological studies, CHMI experiments, and clinical trials (*26, 31, 35, 60–62*). Many infectious disease studies have used the chimeric design of multiple antigens with IEDB resources to select antigenic stretches (*63*). Our chimeric antigens were designed using six different parameters of IEDB resources to select B-cell epitope sequences confidently and were successfully stitched, cloned, expressed and validated. These chimeric antigens were able to get recognized by the human immune response. The epitope mapping showed a heterogeneous IgG and IgM responses that were expected in polymorphic genes and were present across all the peptides selected validating our original design of the antigen. This further predicts that the chimeric antigens may successfully elicit an immune response in humans and can sustain as circulating antibodies.

PfEMP1 proteins are known for their mutually exclusive expression, antigenic variation, and the vast number of endothelial receptors with which they interact. Due to their epigenetic switching and endothelial receptor specificity, targeting any particular PfEMP1 protein is difficult (*40, 41*). Acquired immunity in endemic regions is believed to occur in an orderly, starting from PfEMP1, causing cytoadhesion in vital organs with stronger receptor interaction. As the breadth of immunity increases, parasites are likely forced to express PfEMP1 proteins that interact comparatively weakly and sequester in non-vital organs (*64*). In some cases, multiple rounds of exposure to a sufficient antibody repertoire against PfEMP1 proteins are built to keep the disease asymptomatic (*26, 29, 30*). Sampling all PfEMP1 proteins in their epitope form, irrespective of the subclass, can theoretically protect against cytoadhesion. With antibodies specific to chimeric varB, we observed reduction in the number of adhered RBCs in static assays and a further significant decrease in fluid-based dynamic cytoadherence assays. Our data emphasize the role of flow dynamics in the process of cytoadhesion, as previously shown in other studies and show that anti-PfEMP1 antibodies can inhibit the sequestration at physiological conditions. We also demonstrated the use of a fluid-based cytoadhesion assay in real-time and a quantification method to determine the binding of iRBCs per minute. This method is useful for precisely assessing cytoadhesion and can be used with endothelial cell lines and receptors. The pan anti-PfEMP1 antibody pool when used as a passive immunization in patients with severe malaria will serve as an important additional therapeutic in managing the malaria outcome. The field is yet to comprehensively explore the tissue-tropism seen in malaria infection (*25*). Knowing the exact PfEMP1 domain organization and predicting the organ site of sequestration remains a topic of study with a few reports suggesting a role of sub-type A with DC8 being majorly involved in cerebral malaria through ICAM-1 binding (*24, 50*). While sub-type B is mostly seen in uncomplicated malaria (*65*). These reports vary in their results and an intensive analysis of binding affinities is yet to be done. The knowledge of PfEMP1 sub-type and their endothelial receptor affinities can further equip us with identifying the right pool of anti-PfEMP1 antibodies for passive immunization.

The blood-stage vaccine candidates in phase trials mostly target the merozoite surface proteins to either block entry into the new RBC or function by activating the Fc-mediated innate immune response (*66*). The current leading candidate, RH5, is shown to have neutralizing potential, while candidates such as MSP1 show clearance via activation of the complement pathway or opsonization (*35, 59, 67*). Many of the blood-stage candidates have failed the phase trials for multiple reasons. The antigenic polymorphism and redundancy are main bottlenecks for selecting antigens along with low time-window of invasion and exponentially growing parasite count (*68*). Our study addresses it using combination of multiple peptides from different proteins and shows an additive effect in GIA imparted by the mixture of antibodies specific against chimeric MSP and InvP construct. In a single cycle GIA, the mixture of antibodies works much more efficiently at the concentration of 0.13 mg/ml compared to using antibodies separately. In a natural infection setting, having antibodies against multiple antigens belonging to important functional pathways may outweigh the effect of having a higher antibody titer against just one antigen.

Collectively, we show that, as a proof of principle, multi-protein candidates work more efficiently using multi-epitope than their single counterparts. An effective strategy against malaria will involve a combination of interventions against multiple stages with antimalarial drugs. Targeting the blood stage in a chimeric fashion will help reduce parasitemia, alleviate symptoms, inhibit or reverse cytoadhesion (making iRBCs more susceptible to immune clearance), and help reduce transmission to the next cycle.

## Materials and Methods

### 1. In-silico analysis for selection of antigenic peptides

The eight proteins taken from varB subgroup (PF3D7_1200100, PF3D7_0500100, PF3D7_1300100, PF3D7_0324900, PF3D7_1100100, PF3D7_0937800, PF3D7_0300100, and PF3D7_0400100) and AMA-1, MSP-1, MSP-2 and MSP-3 along with EBA-175, Rh-5, SERA-5, and GLURP were subjected to the analysis in Immune Epitope Database-analysis resource (IEDB-AR). Total of 6 parameters including Bepipred Linear Epitope Prediction 2.0, Chou & Fasman Beta-Turn Prediction, Emini Surface Accessibility Prediction, Karplus & Schulz Flexibility Prediction, Kolaskar & Tongaonkar Antigenicity and Parker Hydrophilicity Prediction were applied to each protein sequence. The scores for each amino acids were aligned and peptide stretch of 25 amino acids having average of score 0.6 and above were considered. For chimeric varB, chimeric MSP (AMA-1, MSP-1, MSP-2 and MSP-3) and chimeric InvP (EBA-175, Rh-5, SERA-5 and GLURP) construct a total of 29 peptides, 20 peptides and 24 peptides respectively were stitched together. The stitched sequence was converted to corresponding DNA sequences using backTransSeq and codon optimized for expression in *E. coli* and cloned into pET28a+ vector containing 6X histidine tag using NdeI and XhoI restriction sites.

### 2. Cell culturing

#### A. Plasmodium falciparum culturing

*P. falciparum* strain 3D7 was cultured according to the previous publication. Briefly the culture was maintained in RPMI1640 base media (Sigma) containing 0.5% AlbuMAX I (Gibco), 25 mM HEPES, 100 μM hypoxanthine (Sigma), 1.77 mM sodium bicarbonate, Glucose, 12.5 μg/ml gentamicin sulfate in 5% CO2 and 37℃ humidified incubator. Parasitemia was kept at 5% with 3% hematocrit (O^+^ human RBCs from healthy donors) and routinely synchronized using 5% sorbitol solution for selecting ring stage. For enriching the late-stage parasites 63% density gradient of percoll was used.

#### B. Mammalian endothelial cell culturing

Human Brain Endothelial Cells (HBEC-5i) were a kind gift from Dr. Hem Chandra Jha (Indian Institute of Technology, Indore). The cells were cultured in DMEM-F12 with L-glutamine, 15mM HEPES, sodium bicarbonate and trace elements (AL139A) HiMedia Pvt Ltd) along with 10% Fetal Bovine Serum (RM9955 HiMedia) and 0.04mg/ml Endothelial Cell Growth factor (ECGS) (02-102 Sigma). Cells were maintained at 70-90% confluency on 0.1% gelatin-coated culture dishes at 5% CO_2_ and 37 ℃ temperature.

### 3. Chimeric protein expression, purification, and mass-spectrometry validation

The three clones (chimeric varB, MSP, and InvP) were separately transformed into a competent *E. coli* BL21DE3 expression system. The culture was grown till the optical density reached 0.6 in 600 nm spectra and induced using 0.5mM of isopropyl-1-thio-β-d-galactopyranoside (IPTG) at 18℃ for chimeric varB and at 25℃ for chimeric MSP and InvP for 8-10 hours. Only the bacterial mass was collected by centrifugation at 4000 RPM and sonicated using sonication buffer (10 mM Tris-Cl pH 7.5, 300 mM NaCl, 10% glycerol, 0.1% TritonX-100, 1X PIC, and 2 mM PMSF) at parameters of 3 sec ON/ 5 sec OFF/ 60% amplitude (Probe sonicator Thomas Scientific). The supernatant was separated by spinning at 12,000 RPM, and Ni-NTA agarose beads (Qiagen 30230) were added to the lysate and incubated for 4 hours at 4℃. The elutions were taken by adding 50 mM, 100 mM, 150 mM, and 300 mM elution buffer (10 mM Tris-Cl pH 7.5, 150 mM NaCl, 10% glycerol, and varying imidazole). The elutions were run on 10% SDS-PAGE gel and stained using Coomassie brilliant blue solution. The elutes containing purified fractions were pooled, dialyzed (10 mM Tris-Cl pH 7.5 and 150 mM NaCl), and concentrated. For validation, the purified proteins were reduced using 10 mM Dithiothreitol, alkylated using 10 mM Iodoacetamide, and trypsinized using sequencing grade modified trypsin (Promega V5113). The samples were desalted and run on Sciex TripleTOF6600. The peptide identification and analysis were performed using MaxQuant.

### 4. ELISA assay

The chimeric antigens at the concentration of 2 μg/ well were coated onto the 96-well plate using 0.05 M carbonate buffer pH 9.6 (0.293 g NaHCO_3_ and 0.159 g Na_2_CO_3_ in 100 ml) at 37℃ for 1 hour. The wells were blocked using 5% Bovine Serum Albumin (BSA) for 1 hour at 37℃. The plasma separated from patient samples was diluted to 1:2000 and incubated for 1 hour at 37℃. The wells were washed five times using 0.01 M PBST pH 7.4, followed by probing with Goat anti-human IgG secondary antibody with HRP conjugated (Invitrogen 62-8420) for 1 hour 37℃. The wells were washed five times and developed using TMB substrate solution (ThermoScientific N301) for 15 minutes in dark conditions. The reaction was quenched using 0.2 M H_2_SO_4_, and the absorbance reading was taken at 450 nm in an Ensight plate reader (Perkin Elmer). A tetanus toxoid positive control and BSA negative control were used for every assay. Each antigen was probed in duplicate and the entire ELISA assay for patient samples were done at least N=2 times.

### 5. Epitope mapping

A 16-array slide containing peptides specific to chimeric varB, MSP, and InvP was designed by PEPperPRINT Gmbh, Germany. The handling and protocol were followed as per the manufacturer’s instructions. The malaria-infected patient samples were used at the dilution of 1:1000 and 1:2000 to detect IgG and IgM-specific responses. The rabbit-generated antibodies were probed at a dilution of 1:500 and images were taken on Agilent SureScan G2600D system. For the analysis of fluorescence intensities, Mapix software was used. The rabbit specific anti-varB antibody was probed N=2 times on two different microarray slides.

### 6. Generation of polyclonal antibodies

The purified chimeric antigens were injected subcutaneously in rabbits (New Zealand White Males aged 12-14 months). Pre-immune sera for each rabbit were collected before injection. Briefly, the antigen preparation involved taking 500 μg of SDS-PAGE separated chimeric varB and MSP proteins (crushed in liquid N2) and 500 μg of chimeric InvP protein in-solution and mixing with an equivalent volume of Freund’s complete adjuvant (Merck F5881). For the subsequent five booster doses, 250 μg of proteins were emulsified with an equivalent volume of Freund’s Incomplete adjuvant (Merck F5506) and injected post 21 days. Bleed was collected after 14 days of booster doses and separated into serum by spinning at 4000 RPM.

### 7. Purification of antibodies

#### A. Chimeric varB

About 3 ml of bleed 3 was diluted in 7 ml of sterile 1X PBS and passed through 0.2 μM filter and incubated with recombinant protein G sepharose 4B beads (Life Technologies 101242) at 4℃ overnight. The beads were collected in a column and washed using 1X PBS and eluted using 100 mM glycine pH 2.5 solution. Elutes were pooled, dialyzed (in 1X PBS), concentrated, and stored in a 25% glycerol solution.

#### B. Chimeric MSP and InvP

Since cysteine amino acids were present in these constructs, we used sulfolink-mediated antibody purification. Around 800 μg of respective antigen was reduced using 25 mM TCEP to form free sulfhydryl groups that bind to the sulfolinkTM coupling resins (Thermoscientific 20401) as the manufacturer’s instructions gave. Once the antigen column is ready, diluted and filtered serum was passed through the column six times and was left for binding overnight at 4℃. The column was washed using 20 mM Tris-Cl pH 7.5, and the antibody was eluted using 100 mM glycine pH 2.5. All elutes were further pooled (separately for MSP and InvP antigen) and dialyzed (in 1X PBS) and concentrated, stored in 25% glycerol. The purification was done N =3 times on the collected bleeds.

### 8. Peptide blocking assay

For validating the purified chimeric MSP and InvP antibodies, an equal amount of late-stage parasite lysate was separated onto 10% SDS-PAGE gel, transferred onto PVDF membrane using a semi-dry transfer method at 25V (Bio-Rad) and the blot was split into two parts. One part was probed with 1:500 dilution of purified antibodies (either chimeric MSP or chimeric InvP), and the other was probed with antibody dilution + 5 μg of purified antigen (either chimeric MSP or InvP) overnight at 4℃. Blots were washed using 1XTBST solution, and anti-rabbit HRP-conjugated secondary (Bio-Rad 1706515) was added at a dilution of 1:4000 for one hour at room temperature. The blots were developed using Clarity Max ECL western blotting substrate (Bio-Rad 1705062) on ImageQuant LAS 4000 system (GE Healthcare).

### 9. Immunofluorescence assay

Late-stage 3D7 parasites were washed and re-suspended in 1X PBS and fixed using 4% paraformaldehyde with 0.0075% glutaraldehyde at 37℃ for 20 min. Permeabalization was done using 0.1% TritonX-100 solution for 10 min at room temperature. Freshly prepared 3% Bovine serum albumin (in 1X PBS) was used for blocking for 1 hour at room temperature. Primary antibodies (purified chimeric antibodies) at concentration 1:500 were added to the suspension, incubated overnight at 4℃/ 8 RPM, and washed using 1X PBS five times. Anti-rabbit Alexa Fluor 488 (Invitrogen A32731) secondary antibodies were added at the dilution 1:500 for 1 hour at room temperature. Imaging was done on the Zeiss Multiphoton microscope.

### 10. Rosette inhibition assay

#### A. Wet mounting

*Plasmodium falciparum* 3D7 synchronized culture was maintained at 3% hematocrit and 5% parasitemia in 10% A+ human serum (heat inactivated at 65℃) for facilitating the rosette formation. 500 μl of culture (after media change and removal of human serum) was incubated with either anti-chimeric varB antibodies or heparin solution. Incubated for 1.5 hours/ shaking platform. 37℃. Suspension was treated with equivalent volume of acridine orange solution (HiMedia TC262) and incubated for 20 mins at room temperature at 8 RPM. 10 μl from the stained suspension was mounted onto the cover slip and imaging done using EVOS microscope (brightfiled and 488 excitation). 5 fields were blindly taken and rosette were defined as a clump of uninfected RBCs surrounding an acridine orange positive infected RBCs. The no. of rosettes were counted in each field manually and plotted using GraphPad prism 7.

#### B. Flow cytometry based

The protocol was adopted from chapter 37 (Malaria Immunology: Targeting the Surface of Infected Erythrocytes, Methods in Molecular Biology. Vol. 2470)(Quintana & Ch’ng, 2022). The staining of RBCs was done using DAPI and DHE. The FACS based sorting of multiplet population was done N=3 times.

### 11. Static cytoadherence inhibition assay

HBEC-5i cells at 70-80% confluency treated with 10 ng/ml of TNF-α for 6 hours before the assay. Parasite culture of 5% parasitemia and 3% hematocrit at late trophozoite and schizont stages were added onto the cells either with range of anti-chimeric varB antibodies or control. Incubated for 1 hours at 37℃ with intermittent shaking. Unbound RBCs were removed with 3 times sterile DPBS washes. Cells were fixed using 1% glutaraldehyde solution and stained using 10% geimsa for 20 mins at room temperature. Imaging done on Leica DM750 with Flexacam coloured camera. From each experimental conditions 15-20 random fields were taken and number of HBEC-5i cells and iRBCs were counted. The data is plotted as a ratio of iRBCs present to the HBEC-5i cells visible in the field.

### 12. Fluidic cytoadherence inhibition assay

HBEC-5i cells (cell count = 180,000 per lane) treated with TNF-α (10 ng/ml) resuspended in 25 μl of media were seeded onto a lane of the chip (BeOnChip-Be flow Microfluidic chip, OOC-FLOW-01, Fluigent Smart Microfluidics) precoated with 0.1% gelatin. Cells were supplemented with 200ul of media, allowed to adhere for 24 hours, and treated with TNF-α for 6 hours before the assay. Parasite culture was diluted in media to achieve 0.75% hematocrit (higher hematocrit makes the visualization on camera difficult for live tracking of adhered iRBCs). It was treated with anti-chimeric varB antibodies in the range of 10 μg and 20 μg per 2 ml for 10 mins at 37℃ before passing from the chip. The culture was passed at the flow pressure of 25mbar and a 1-minute video was taken every 2-3 minutes during the passing of the culture using Leica DM750 with Flexacam.

### 13. Quantification of adhered iRBCs

To quantify the number of adhered RBCs, an in-house detection and analysis pipeline was developed and is briefly described here. Bright-field microscopy images of RBCs in the microfluidic cell show a high contrast in the central region similar to that seen in previous work (*69*). We used image filtering by a Laplacian of Gaussian (LoG) to enhance local contrast, followed by image binarization and centroid detection. The resulting RBC coordinates detections. that appear in the same location in 5 adjacent frames are considered as stationary cells and retained for further processing. Since these regions could correspond to either adhered RBCs or background noise, a second selection step is implemented, where a shallow convolutional classification network is utilized to select “in-focus” RBCs while discarding the rest. The network was trained on manually selected in-focus RBCs as the positive class, with spurious detections classified as background. To further enhance the detection quality, a probability threshold of 0.93 is applied. This threshold ensures a higher confidence level in selecting RBCs, potentially reducing false positives and improving result accuracy. The classifier achieves an validation accuracy rate of 95.75% on validation data. (Supplementary Figure 3B and C).

### 14. Growth Inhibition assay

Parasites at 1% parasitemia (synchronized ring stage) maintained at 3% hematocrit were seeded onto a 96-well plate with an indicated range of purified anti-chimeric MSP and anti-chimeric InvP antibodies (either alone or in addition). This was done over two cycles of invasion (media change with the addition of antibodies). At the end of the second invasion, the plate was processed for lysis (100 mM Tris-Cl pH 7.5, 25 mM EDTA, 0.05% Saponin, 0.5% TritonX-100, and 1X sybr green dye) for 40 mins in RT/dark conditions. The fluorescence readings were taken at 497 nm excitation and 520 nm emissions. Fluorescence intensity was normalized to the readings of uninfected RBCs and plotted using GraphPad Prism 5. The GIA was done using active and inactive bleeds (data not shown) and purified antibodies in total N=9 times.

### 15. RNA sequencing of cytoadhered cells

The parasite line 3D7 was allowed to cytoadhere to HBEC-5i cells as per mentioned in materials and method section 11. The bound fraction was collected by dislodging them using stronger washes. The collected cells were passed through a cellulose column packed in a syringe to remove possible contamination of HBEC-5i cells. After enrichment of iRBC, they were subjected to lysis using TRIzol (Biorad) and further RNA isolation was done using phenol-chloroform extraction method. The RNA quantity and quality was determined by Qubit fluorometer (Invitrogen) and Agilent Tapestation respectively before using for RNA sequencing on Illumina platform. The RNA library preparation was done using NEBNext Ultra II RNA library prep kit (NEB E7770S) as per manufacturer’s instruction. The sequencing was done using 75 cycle high output paired end format.

### 16. Data analysis of RNA sequencing

The quality of the sequencing was determined using FASTQC and adapter sequences were trimmed using Trimgalore. The alignment to *Plasmodium falciparum* genome was done using HiSat2 and further TPM generation was done using StringTie analysis. The TPM values for PfEMP1 genes were separated and plotted using Graph Pad Prism 7.

## Ethics Statement

We are thankful to the Department of Medicine, S.P. Medical College Bikaner for the collection of patient serum samples (No. F. 29 (Acad)SPMC/2020/3151). Human RBCs used in this study were obtained from the KEM Blood Bank (Pune, India) as blood from anonymized donors. Approval to use this material for *P. falciparum in vitro* culture has been granted by the Institutional Biosafety Committee of Indian Institute of Science Education and Research Pune (BT/BS/17/582/2014-PID). The use of rabbits in this study for immunization (IISER/IAEC/2017-01/008) was reviewed and approved by Indian Institute of Science Education and Research (IISER)-Pune Animal House Facility (IISER: Reg No. 1496/GO/ReBi/S/11/CPCSEA). The approval is as per the guidelines issued by Committee for the Purpose of Control and Supervision of Experiments on Animals (CPCSEA), Govt. of India.

## Acknowledgements

We are grateful to all the volunteers and guardians of children who have consented to provide blood samples for this study. We are very grateful to Dr. Hem Chandra Jha (Indian Institute of Technology, Indore, India) for kindly gifting us with the HBEC-5i cell line. The illustrations presented in the manuscript are created using BioRender.com. This work was supported by DBT-Infectious Disease Biology division (BT/PR41408/MED/29/1547/2020) and MoE-STARS-2/2023-0249 from the Government of India to KK. The funders had no role in study design, data analysis, decision to publish, or preparation of the manuscript. We are thankful to the NGS facility, IISER Pune for the sequencing support, IISER Mass-spectrometry facility for their proteomics support and IISER FACS facility for their support in flow cytometry experiments.

## Authors’ contributions

BD designed, performed experiments, and analyzed data. DK and CA helped with the quantification of bound RBCs in microfluidic assays and associated experimental design. SK provided patient serum samples and helped with the analysis and experiment design. BD and KK wrote the manuscript. KK planned, coordinated, and supervised the project. All authors read and approved the final manuscript.

## Supplementary Information

**Supplementary Figure 1. Expression, purification and validation profile of chimeric antigens. (A)** Transcript expression of all PfEMP1 genes in lab grown 3D7 strain (adapted from Rawat et al., 2021) with genes used in chimeric varB marked in red. **(B)** Western blot probed using anti-his antibodies showing expression pattern of chimeric varB (left), MSP (middle) and InvP (right) antigens after induction in BL21DE3 strain. **(C)** Mass-spectrometry based validation of chimeric varB, MSP and InvP construct showing unique peptide counts detected. **(D)** Immuniprecipitation followed by Mass-spectrometry of ptient serum samples showing presence of antibodies against proteins selected for construction of chimeric antigen,

**Supplementary Figure 2. Peptides used in construction of chimeric antigens show diverse IgM response in patient samples. (A)** Chimeric varB construct **(B)** chimeric MSP construct and (**C)** chimeric InvP construct.

**Supplementary Figure 3. Rabbit generated anti-chimeric varB IgG antibodies are specific and show inhibition of cytoadhesion under fluidic conditions. (A)** IgG response of rabbit generated anti-chimeric varB antibodies against each peptide used in construction of chimeric varB antigen. **(B)** Detection of intensity peaks corresponding to potential RBCs obtained using DICOT segmentation workflow (*58*) **(C**) that were classified as “in-focus” RBCs using a custom CNN classifier (described in the Materials and Methods section).

**Supplementary Table 1.** A list of peptides along with their protein of origin, location and antigenicity scores of chimeric varB (sheet 1), MSP (sheet 2) and InvP (sheet 3).

**Supplementary Video 1**. Cytoadhesion shown by iRBCs on primed HBEC-5i cells under 25mbar fluidic pressure. Video taken during the culture perfusion showing bound RBCs not moving across time.

**Supplementary Video 2.** Inhibition of cytoadhesion upon addition of anti-chimeric varB antibodies. **(A)** control condition **(B)** Inhibition under 0.02 mg/ml anti-chimeric varB antibody concentration **(C)** Inhibition under 0.04 mg/ml anti-chimeric varB antibody concentration.

